# A physically inspired approach to coarse-graining transcriptomes reveals the dynamics of aging

**DOI:** 10.1101/2024.03.13.584889

**Authors:** Tao Li, Madhav Mani

## Abstract

Single-cell RNA sequencing has enabled the study of aging at a molecular scale. While substantial progress has been made in measuring age-related gene expression, the underlying patterns and mechanisms of aging transcriptomes remain poorly understood. To address this gap, we propose a physics-inspired, data-analysis approach to extract additional insights from single-cell RNA sequencing data. By considering the genome as a many-body interacting system, we leverage central idea of the Renormalization Group to construct an approach to hierarchically describe aging across a spectrum of scales for the gene expresion. This framework provides a quantitative language to study the multiscale patterns of aging transcriptomes. Overall, our study demonstrates the value of leveraging theoretical physics concepts like the Renormalization Group to gain new biological insights from complex high-dimensional single-cell data.

## Introduction

Recent advances in RNA sequencing technologies [1] now allow the high-throughput quantification of genome-wide gene expression in organisms at single-cell resolution [2]. The Tabula Muris dataset [3] which profiles mouse transcriptomes across different age groups provides a starting point for studying the dynamics of aging. In particular, these data include gene expressions across the life of mice, which permits the high-resolution genome-wide understanding of aging dynamics. Performing linear regression between individual gene expression and age identifies the most positively and negatively age-correlated genes [3]. However, single-gene level analyses aren’t sufficient to wholistically capture the dynamics of aging as manifest in transcriptomes. Inspired by the renormalization group (RG), we propose a physics-inspired, data-analysis approach to integrate across different scales of gene expression and construct a collective, multi-gene, description of aging [4, 5].

Meshulam et al. [5] outlined an approach to data analysis inspired by the Renormalization Group [6, 7]. Summarizing, an appealing analogy to real space and momentum space renormalization group is presented for analyzing high-dimensional data: 1) Analyze the evolution of correlations as variables are iteratively paired based on correlation coefficients; 2) Study how correlations change as progressively smaller linear modes of variation are retained. Though Meshulam and Bradde et al. analyzed imaging data of a dynamical population of neurons [5] and market return data [4] and searched for the criticality, this approach can be further pursued in various high dimensional data contexts. Here, we attempt to pursue this in the context of a single-cell RNA sequencing-based assay of aging mouse tissues. This approach juxtaposes the usual approach to such transcriptomic data, which typically involves the projection of high-dimensional data into a lower-dimensional space wherein an attempt to identify clusters is made [8], as noted in [9, 10]. However, this assumes an ideal scale exists at which to render the system understandable. This is contrary to an investigation of physical systems where system features, such as correlations, are studied as a function scales in the system [11].

Implementing some of the salient ideas from RG to biological contexts poses a challenge. In most “physics” scenarios where RG is used, observables of a system are studied and combined according to their geographical location within a system. However, as is the central point of Meshulam et al., [5], in traditional physical systems, especially the system on a grid (e.g. spin net), space is merely a convenient parametrization for interaction strengths which are local. Said another way, interactions, and their relative strengths, are the fundamental objects of interest. We apply Meshulam et al.’s RG-inspired framework, which uses correlation as a proxy for locality, to extract multiscale insights into aging transcriptomes.

Additionally, most high dimensional datasets are severely undersampled [12] and thus more simple-minded approaches to data-analysis that leverage covariance and correlation statistics may provide more robust and interpretable results compared to approaches relying on detailed modeling of the underlying joint probability distribution or underlying mechanics. However, some argue that linear techniques like PCA are unsuitable for real-world data, since the eigenvalue spectrum is typically continuous rather than exhibiting clear spectral gaps [4]. Thus, the traditional way for identification of latent dimensions is not always feasible. However, as Bradde et al. discuss [4], this continuity does not preclude the value of PCA. Unlike conventional PCA which seeks a single low-dimensional latent representation, combining PCA with coarse-graining enables hierarchical characterization of the data. Avoiding imposition of discrete scales, this physics-inspired approach can provide multi-scale insights even for highly undersampled and continuous eigenvalue spectrum data where traditional PCA falls short.

We apply the aforementioned approaches to analyze a recent single-cell transcriptomic dataset profiling mouse tissues across ages. Studying aging is well-suited for these physics-inspired methods since the underlying trends are expected to be continuous and quantitative across time. Rather than identifying discrete novelties like new cell types or reducing the data to a low-dimensional representation, the goal is to connect different scales of the system using RG-inspired coarse-graining.Specifically, this multiscale analysis aims to incorporate features from various scales of resolution into downstream analyses.

In Sec 1, we describe the transcriptomic data and normalization approach. Then, two formulations of coarse-graining methods are proposed. Next, in Sec 2, we demonstrate the basic results at different scales from both methods using data from a single tissue type. Leveraging these outputs, we then present a spectral view of aging by comparing to null models, focusing on two key aspects: the emergence of block structure in the correlation matrix, and the normality of individual genes quantified by 4th order moments and Anderson-Darling statistics. This spectral characterization reveals a shift between single gene Gaussianity and block structure in correlation of entire genome, which is not detectable without coarse-graining analysis.

### 1 Transcriptomic Data and Coarse-graining Method

We leveraged single-cell RNA sequencing data from the Tabula Muris Senis project [3] (Fig. 1), which profiled 21 mice using droplet-based sequencing technologies. This dataset contains count matrices for individual cells from various organs across the entire lifespan(Fig. 1. Fig.1(b) shows Uniform Manifold Approximation and Projection (UMAP) [13] plots of the normalized data (described below), with clear separation between tissue types. Each UMAP plot corresponds to one gender and age group, providing an overview of the transcriptomic landscapes.

**Fig 1.**
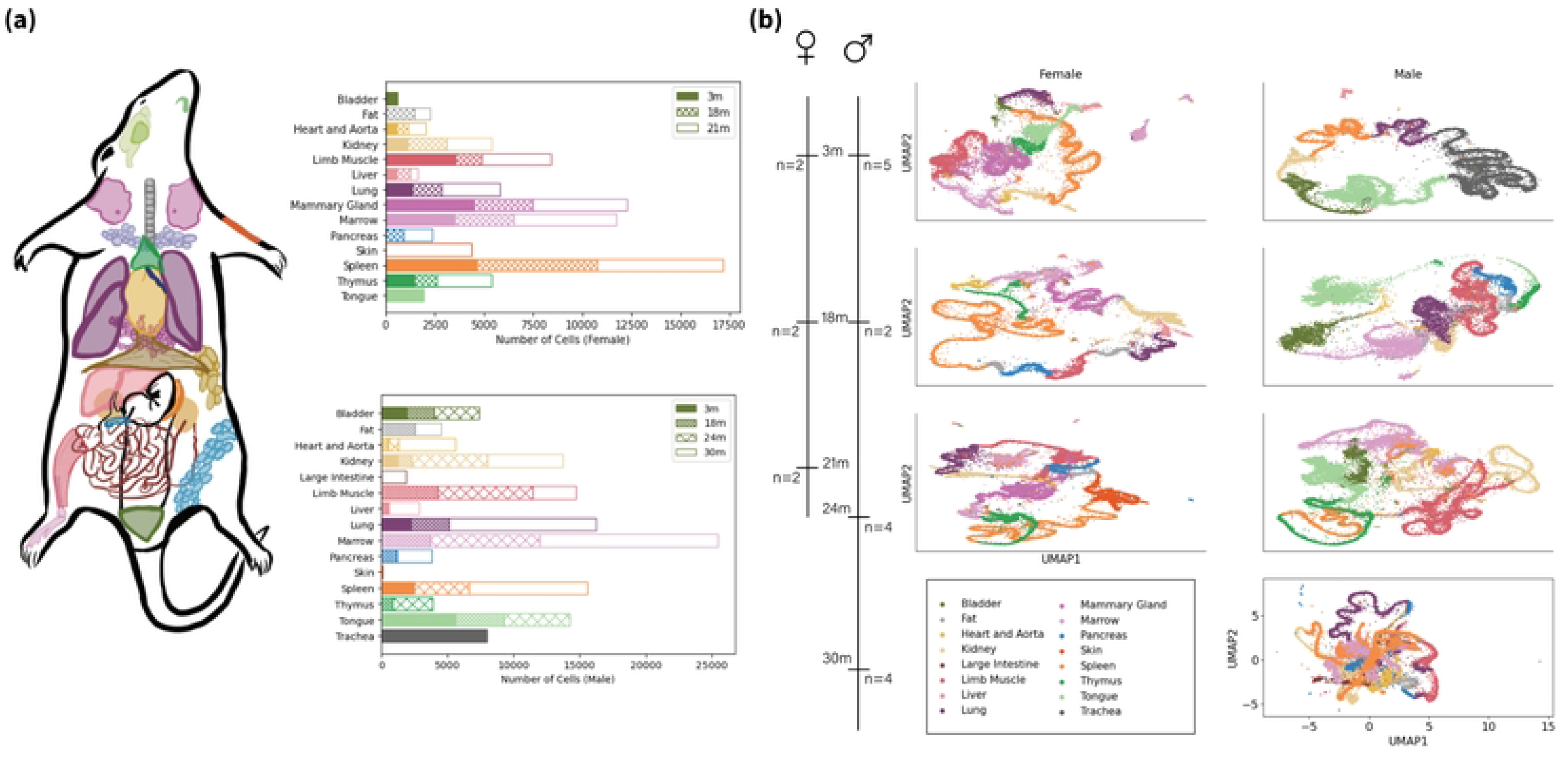
Overview of the single-cell transcriptome data. **(a):** Color-coded tissues that are sampled from Mus musculus by the Droplet method and the number of counts for each combination of age and tissue; **(b):** The distribution of sex and age, the corresponding UMAP visualization of the raw data.

As shown in Fig. 1, spleen, mammary gland, bone marrow, and limb muscle tissues were sequenced at high depth in this dataset. However, not all cell types from these 4 tissues were profiled across the full age range of 3 months to 21 months. Therefore, we focus our analysis on 4 key cell types, including 2 from the spleen: B cells, T cells and mammary gland T cells and limb muscle cells. Restricting to these subsets, the entire timeline of young to old mice can be covered.

#### Normalization

The raw gene expression counts exhibited heteroscedasticity, with gene- and cell-specific variability [14]. To account for this, we normalized the count matrices using the analytical Pearson residual method [15]. This approach assumes negative binomial distributed counts, with gene- and cell-specific mean and variance. The method fits these variations, enabling inference of normalized z-scores for each count by approximating the mean and variance (see Methods). Normalization restores the counts of genes to an equal footing (SI 3 Fig. 6), allowing us to focus solely on correlations for subsequent coarse-graining analysis.

#### Coarse-Grain Approach in Real Space

We apply the coarse-graining approach inspired by Meshulam et al. [5] to the Tabula Muris Senis data. Conventionally, coarse-graining is based on locality in physical space, with local interactions between microscopic variables, as pointed out in famous spin net example [6]. Following Kadanoff’s block-spin principle [6], nearby variables are aggregated into coarse-grained variables, presuming local variables must have strong correlation. Contrasting such classical uses of these ideas, correlation strength need not be solely determined by physical proximity. In the case of gene expression data, the strength of correlations between two genes isn’t related to their proximity in 3D physical space, precluding spatial locality-guided coarse-graining. However, each gene’s expression state can be viewed as a random variable. Sequencing, a measurement of the collective state of gene expression in single cells, provides samples from distinct configuration of their joint probability distribution. The strength of gene-gene correlations can thus be estimated using sample correlation coefficients. Then, inspired by block spinning, we use correlation strength itself to define “neighborhoods” and study the evolution of the correlation structure through coarse-graining. Specifically, we calculated the gene-gene correlation matrix and greedily paired highly correlated genes. Metagenes were defined by averaging paired gene expression. This pairing is iterated until only one coarse-grained gene is left. Fig. 2 illustrates the real-space coarse-graining procedure. Each ellipse is a collection of an ensemble of cells with the normalized expression of **one** specific gene. At the original scale (step 0), colored ellipses show example gene expression distributions, with red being, relatively, highly expressed gene and green otherwise. Then, highly correlated genes are paired based on ranked correlation coefficients. Each gene pair is averaged to define a metagene for the first coarse-graining iteration. This pairing and averaging procedure is repeated on the metagenes to further coarse-grain the data. While the mean expression remains zero through coarse-graining, the variance fluctuates. To impose equal footing assumption to metagenes, the variance of metagenes is set to 1 at each iteration. This enforces coherent units across resolutions and enables analyzing the evolution of correlation structure.

**Fig 2.**
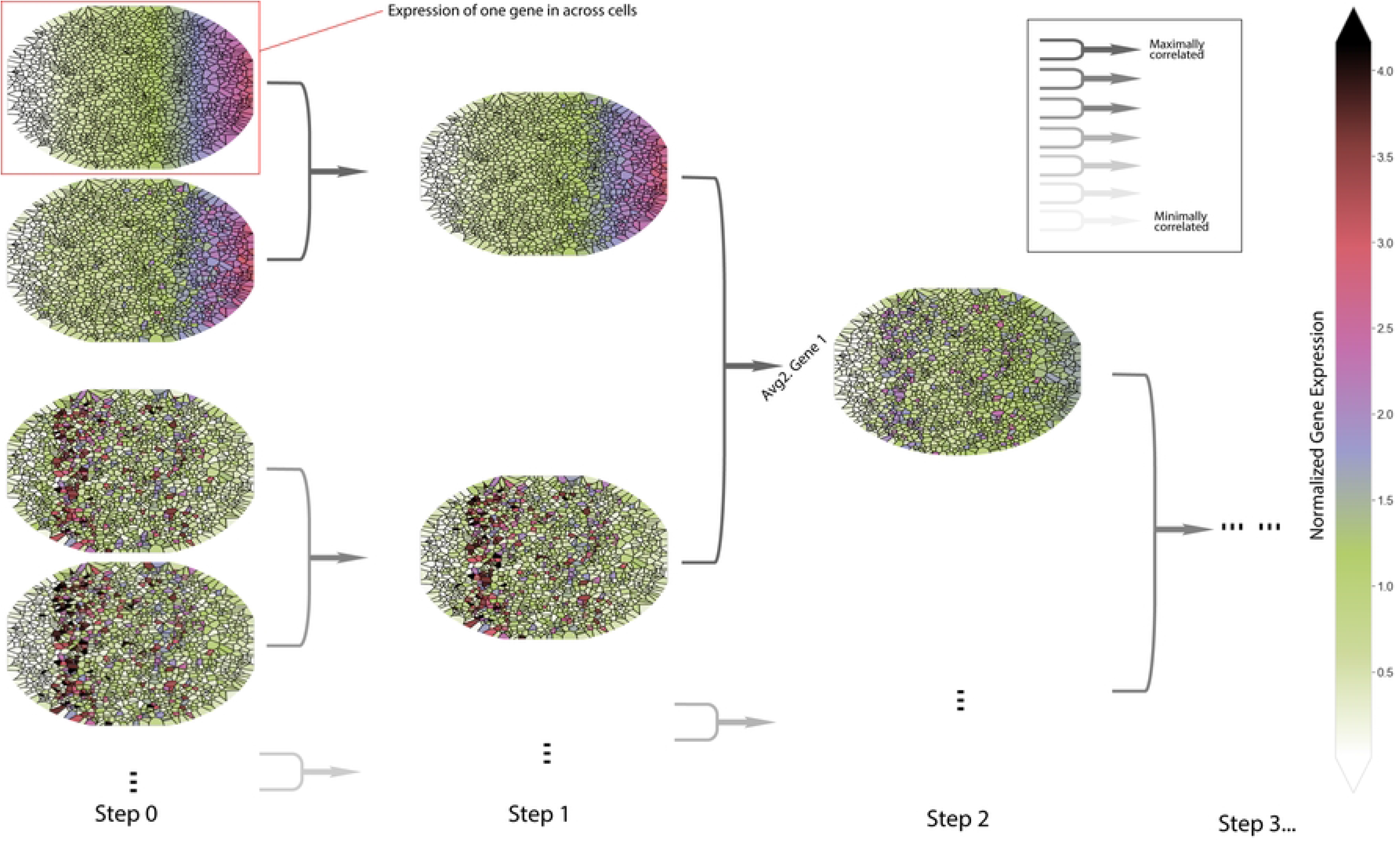
Real-Space Coarse-Graining. All ellipsis represent the same system, such as a tissue, an organ, or several organs. Small areas segmented by solid lines represent single cells. Every gene has an expression distribution across all cells as per ellipse, and the color codes for the expression level. At step 0, the data is the normalized single-cell atlas. At each following step, maximally correlated variables get paired to produce a coarse-grained metagene then the second maximally correlated pairs. The procedure is iterated until only one gene is left where we cannot coarse grain further.

Formally, we start with *N* genes 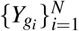 and calculate all correlation coefficients for every possible pairs *ρ*_*i, j*_(*i* ≠ *j*). We then greedily search for the maximally correlated pair, denoted genes 1 and 2. A coarse-grained variable (metagene) is constructed as follows

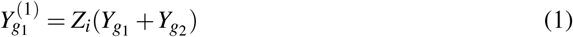

where *Z*_*i*_ normalizes each metagene to variance 1. This pairing is repeated for the next most correlated pair, gradually reducing the N original genes to ⌊*N/*2⌋ metagenes describing a depth 1 coarse-grained system. Iterating this process generates a flow of coarse-grained systems with ⌊*N/*2^*k*^⌋ variables at depth *k*.

#### Coarse-Graining Approch in Momentum Space

Momentum-space coarse-graining in experimental data starts with PCA and can be achieved by progressively projecting out principal components (PCs) explaining low variance, corresponding to local fluctuations [4]. By doing so, the global structure expected at a coarser scale will be pronounced. Formally, if we start with *N* normalized variables 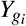, with *i* = 1, 2, …, *N*. These variables will have zero means and be restored to unit variance according to our normalization scheme. We construct the covariance matrix C and performed eigenvalue decomposition,

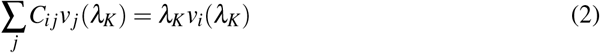

where *λ* is the eigenvalue and *v*(*λ* ) is the associated eigenvector, sorted from largest to smallest. Momentum-space coarse-graining projects out low variance modes containing minimal structure information:

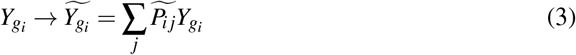

where 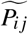 is the projection operation, assuming there are *K*^*^ modes left (not averaged out),

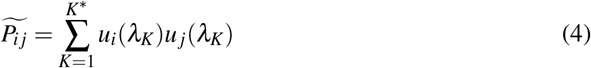

As in standard renormalization group, we tracked the full joint distribution evolution to follow the results.

In physics, renormalization group flows are analyzed by tracking the evolution of Hamiltonians and correlations across scales [6, 7]. For experimental data, studying the full and exact Hamiltonian is generally infeasible. However, Hamiltonians directly relate to the joint probability distribution of all variables. Therefore, for real data, an appealing analogy to track is the joint distribution itself (or marginal) and the correlation structure across scales. As in Monte Carlo simulation studies [16], we focus on marginal distributions of individual coarse-grained variables and their correlation matrices to characterize the full joint distribution under coarse-graining. This physics-inspired analysis of marginal distributions and correlations enables extracting insights into the multiscale structure of real data.

## 2 Results

### 2.1 Inferred correlation structure and coarse-grained joint probability distribution as a function of scale

We apply coarse-graining analysis to an example single-cell RNA sequencing data, 3-month old spleen B cells from two female mice (∼3000 cells, ∼6000 genes), as noted in Sec 1. Prior to presenting our results from coarse-graining, we first perform a null analysis, using marginally resampled data, in order to establish the expected baseline behavior in the absence of genetic correlations(See Methods). To be more specific, the single-cell sequencing data is shuffled for each gene across cells. Null data analysis reveals that (Sec S3 Appendix & Fig. Null Analysis), lacking the correlations, both coarse-graining methods exhibit a convergence to Gaussian distribution. In the absence of correlations, viewed as random variables, metagenes in real-space coarse-graining are indeed expected to converge to a Gaussian distribution, as predicted by Central Limit Theorem. On the other hand, equal variance random modes in marginally resampled data lead to rapid convergence to Gaussian distribution when these modes are sequentially discarded. This null analysis provides an essential baseline to further identify statistically significant non-random features in real data. For example, statistics from normality test can be used to quantify the flow evolution. Additionally, correlation block structure is defined as groupings of variables that have strong inner-group correlations while weak intra-group correlations. Metagenes may exhibit emergent clustering which is not present at single-gene resolutions, as the whole system is coarsened and mesoscale structure is revealed. We measure the strength correlation structure by the spectral gap of correlation matrix, the maximum gap of sorted eigenvalues.

Fig. 3 shows the coarse-graining analysis of Spleen B cells. Fig. 3(a) indicates the portion of the data used. Fig 3 (b) and (c) show the real-space and momentum-space coarse-graining flows, respectively, quantified by the evolution of joint distribution of (meta)genes (b.ii) & (c.ii), correlation block structure or spectral gap (b.i) & (b.iii), and 4th order moments (c.iii). The choice for quantification of the flow is based on the established baseline behavior, Gaussian, which is presented by the black solid curves in (b.ii) and (c.ii,iii).

**Fig 3.**
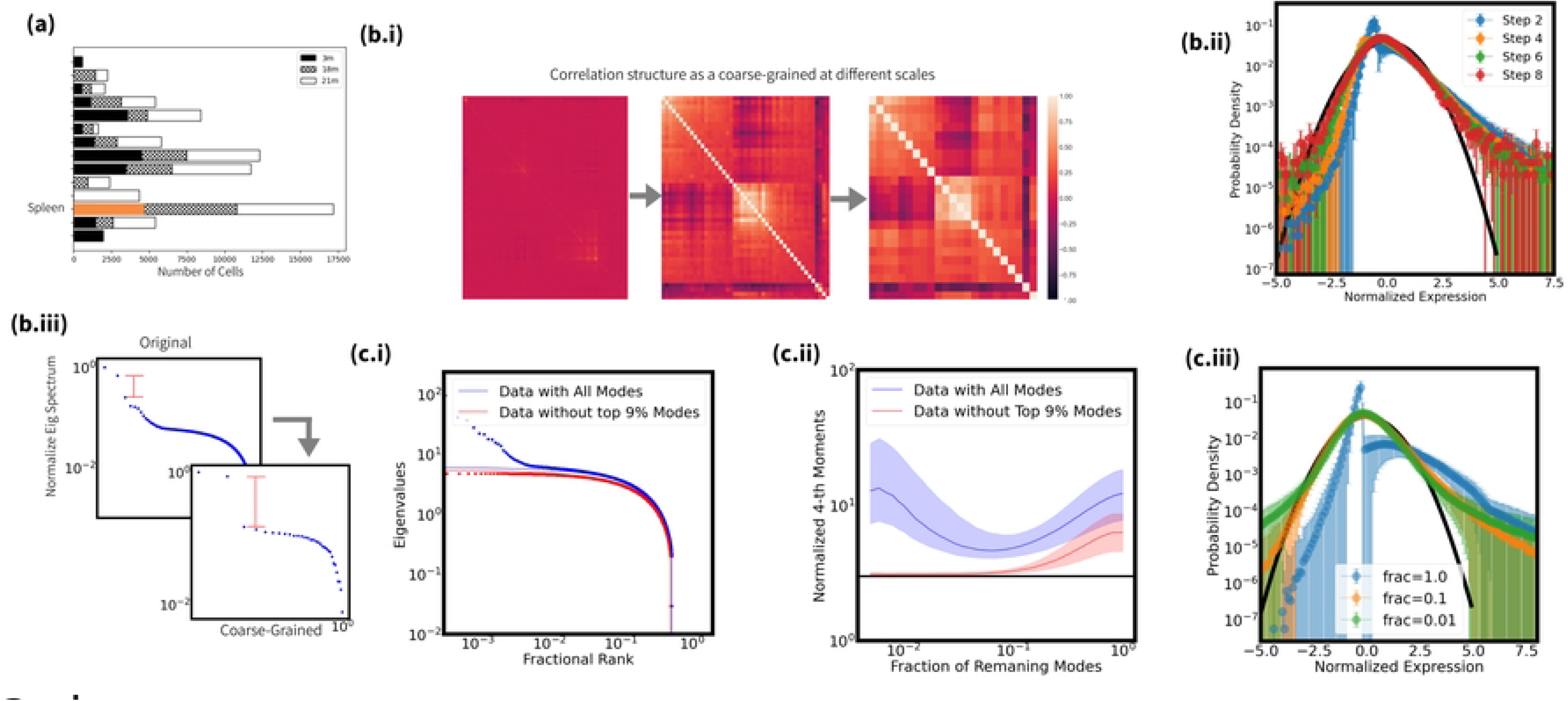
Coarse-Graining Analysis of **Female Spleen B-Cell at 3 month.(a):** The barplot of transcriptome of interest from Figure 1; **(b):** Real-Space Coarse-Graining; **(b.i):** The evolution of correlation matrices, indices (rows and columns) are ranked by the correlation strength within clusters. Block correlation structure is more and more pronounced as we coarse grain; **(b.ii):** The evolution of the probability density of gene expressions for individual (meta)genes as we coarse grain. Solid black curve represents a standard Gaussian distribution;**(b.iii):** The spectral gap becomes increasingly large as we coarse grain.**(c):** Momentum-Space Coarse-Graining; **(c.i):** The eigenvalue spectrum (scatter plot) of the correlation matrix for the transcriptomes and the Marchenko-Pastur distribution (line) excluding 0%(blue) and 9%(red) largest eigenvalues; **(c.ii):** The evolution of the normalized fourth moments of individual genes against the fraction of remaining top modes are plotted with lines as well as one quantile above and below, excluding 0%(blue) and 9%(red) largest eigenvalues. **(c.iii:)** The evolution of the probability density of individual variables when there are only 100% (blue), 70%(orange), 40%(green) and 10%(red) largest eigenvalues left. Black line is proportional to a Gaussian distribution;

Under real-space coarse-graining, the joint distribution remains non-Gaussian at all scales (Fig. 3 (b.ii)), converging to a distribution with fat tails, while correlation block structure becomes increasingly pronounced (Fig. 3 (b.i,iii)). The momentum-space coarse-grained distribution also evolves non-trivially, stabilizing to a non-Gaussian shape with fat tails after removing 90% of low variance modes (Fig. 3 (c.iii)). The discarded modes alone fit a Marchenko-Pastur distribution well (Fig. 3 (c.i) red curve), indicating the removed modes resemble noise and may not be explanatory in this example cell type. However, including the top few high variance modes reveals non-Gaussian evolution. These detectable deviations from the Gaussian baseline will be then leveraged to quantify the coarse-graining flow.

In brief summary, both coarse-graining approaches uncovered non-trivial multiscale structure inaccessible from the original scale alone. Unlike dimensionality reduction methods seeking a single representative scale, coarse-graining establishes a spectral characterization as a function of scales.

### 2.2 Coarse-graining reveals multiscale aging-related changes in transcriptomic structure

Going beyond tracking the evolution of the correlation structure and joint probability as a function of scale, we wish to investigate how the multiscale description of gene expression revealed by coarse-graining evolves with aging. Juxtaposing most prior work, focused on identifying individual genes with age-correlated expression, we propose that a coarse-graining analysis, incoporating information across scales, may provide additional insights into aging.

We applied both real-space and momentum-space coarse-graining to spleen B cells from female mice at 3 ages: 3 months (∼3000 cells), 18 months (∼4000 cells), and 21 months (∼4000 cells). These cells were abundant across all samples, enabling an analysis of aging . Fig. 4 shows the results. At the original single-gene scale, age-related differences in correlation structure are not immediately apparent with minimal spectral gaps. However, after moderate coarse-graining (3 iterations), younger mice (3 and 18 months) exhibited stronger 2-block correlation structure compared to the oldest mice (Fig. 4(b.i)). This difference became more pronounced with further coarse-graining (Fig. 4(b.i)), with the spectral gap being larger for younger mice (Fig. 4(b.ii)). On the contrary, the oldest group shows a more continuous eigenvalue spectrum, indicating weaker block structure (Fig. 4(b.ii)).

**Fig 4.**
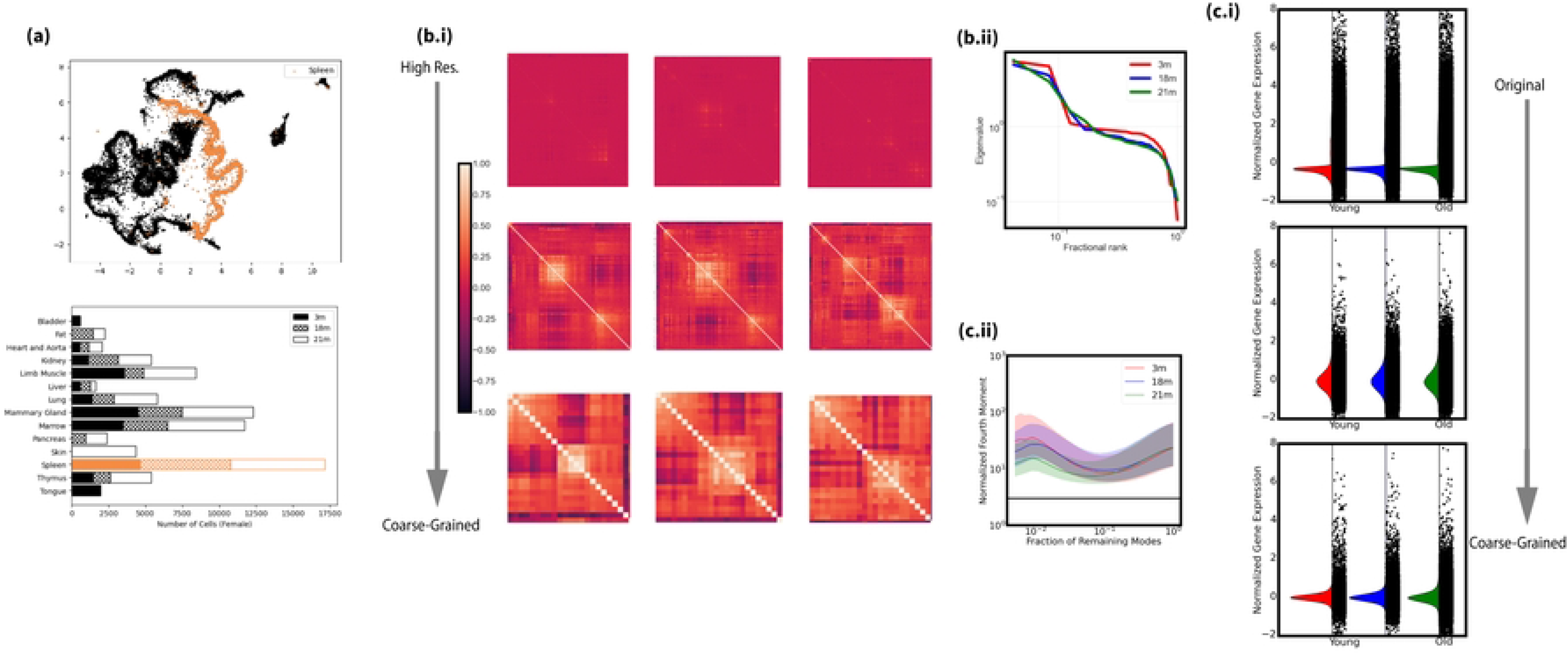
Spetral View of aging (Female Spleen B Cell):**(a):** The UMAP and barplot of the transcriptome of interest from Figure 1; **(b.i):** The spectral view of aging dynamics in terms of correlation structure in real-space coarse-graining. From top to bottom is original scale, coarse-grained for 3 times and c arse-grained for 6 times while from left to the right is from young to old. **(b.ii):** The aging dynamics in the most coarse-grained scale, represented by the eigenvalue spectrum of correlation matrices; **(c.i):** The distribution of genomes at different scales obtained by momentum-space coarse-graining. The distribution is presented by the median distribution as we have done in the previous section. The 3-month, 18-month and 21-month groups are colored in red, blue and green respectively and from top to bottom is from original scale to the most coarse-grained scale. **(c.ii):** The evolution of standardized fourth-order moments (kurtosis), used as an example of quantifying coarse-graining flow.

Momentum-space coarse-graining revealed similar trends. While 4th order moments were indistinguishable at the single-cell scale, differences emerged after coarse-graining, with younger mice showing larger values indicating greater non-normality (Fig. 4(c.i)). Further projection of low variance modes amplified these distinctions (Fig. 4(c.i)).

Summarizing, coarse-graining revealed multiscale aging-related changes in transcriptomic structure inaccessible from original single-cell data alone. For the example cell type studied, younger mice exhibited stronger modular correlation structure and non-normality at emergent coarse-grained scales compared to old mice. The goal of coarse-raining isn’t to find the best scale. Rather, the information encoded across scales composes the spectral analysis.

### 2.3 Multiscale analysis on additional cell types reveals multiscale aging dynamics

We have demonstrated the proposed framework for analyzing high-dimensional gene expression data, using female spleen B cell as an example. Going further, we expand the analysis to a wider set of cell types, which consists of spleen B cell, spleen T cell, mammary T cell and muscle cell. Continuing the quantification in previous section, we quantify aging dynamics using the Anderson-Darling normality statistics and spectral gaps. Larger spectral gaps indicate independent correlation blocks, while higher normality test statistics correspond to lower normality (Fig. 5(a,b)). Spectral gaps were normalized by the largest eigenvalue for comparison, and normality tests were adjusted for sample size (see Methods) [17].

**Fig 5.**
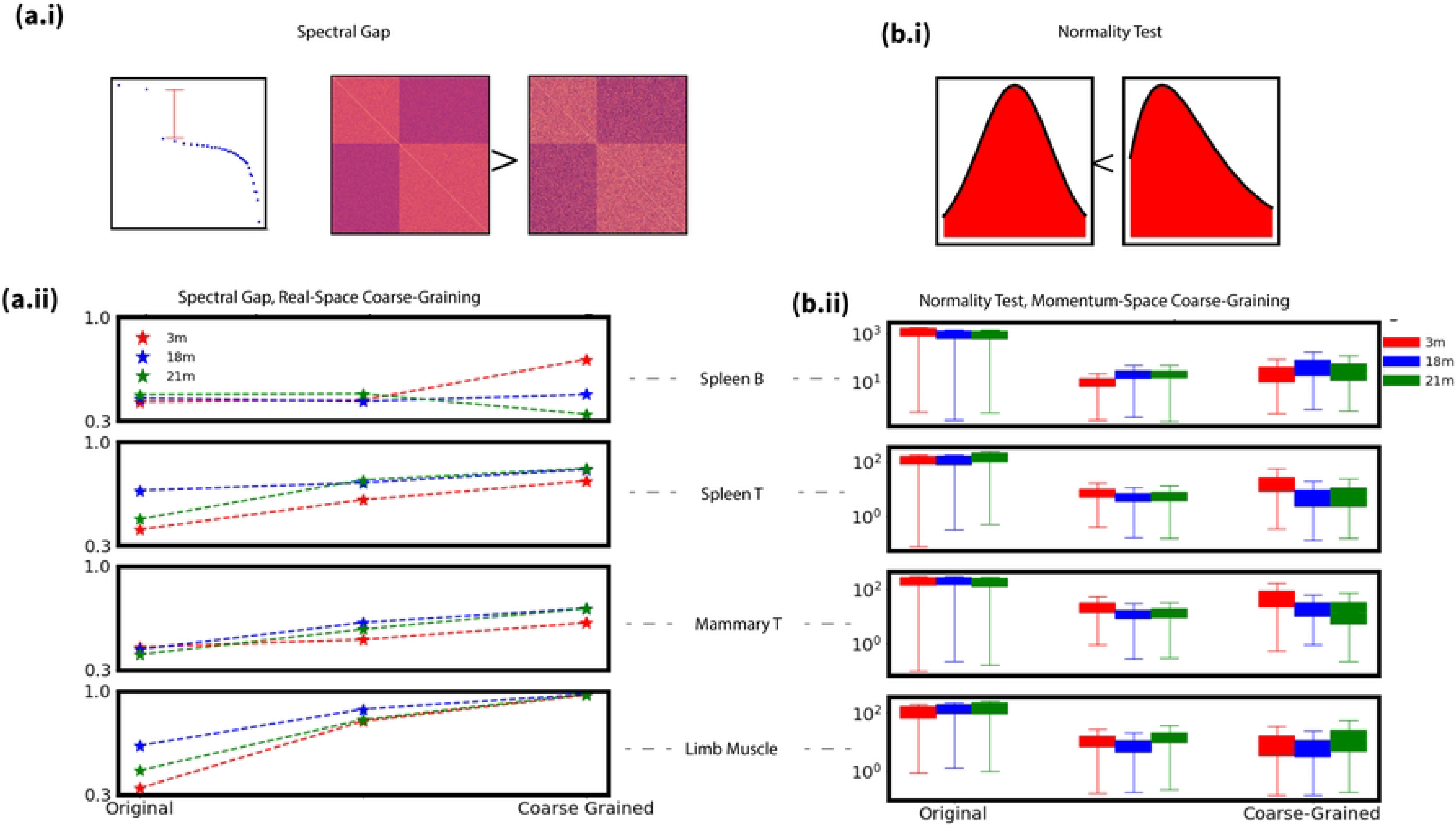
Example of Spetral Modeling of aging. **(a.i):** An intuitive demonstration of spectral gap, visually similar block structure will have a higher spectral gap if the block structures are ‘more uncorrelated’. **(a.ii):** The spectral gap of correlations for real-space coarse-graining. **(cb.i):** The usage of Anderson-Darling statistic, a measurement of normality. **(b.ii):** Distribution of AD statistics of each single gene for momentum-space coarse-graining.

Aging in four cell types are then demonstrated(Fig. 5 (a,b)). At the single-gene scale, aging trends are unclear for some cell types. However, coarse-graining reveal additional insights. For example, in spleen B cells, coarse-graining shows a decreased spectral gap with aging, indicating weaker block structure (Fig. 5(a)). On the other hand, it is also possible that different age groups exhibit coarse-grained features, such as limb muscle cell.

Relying on these two metrics, we construct a quantitative spectral modeling of aging dynamics for four different cell types (Fig 5 (a.ii) & (b.ii)). At the original scale, it could be difficult to specify aging dynamics in terms of the degree of normality. Or in terms of the spectral gap, the original scale does not well indicate that aging affects the correlation structure, as shown in spleen B cell or Mammary T cell. As one should expect, different cell types demonstrate distinct aging dynamics. Spleen and mammary T cells show opposite trends to spleen B cells, with increasing spectral gaps and increasing normality with aging (Fig. 5). Limb muscle cells exhibited non-monotonic changes, with peak block structure and normality at maturity.

Despite the above cell type-specific differences, we also identify an intriguing universal signature - a shift between spectral gaps and the normality of single genes during aging. More specifically, increased normality statistics (less Gaussianity) is accompanied by decreased spectral gaps (weaker block structure) with aging across multiple cell types, and vice versa. For example, in spleen B cells, genes became less normally distributed across cells with age based on the Anderson-Darling test, while concurrently, the coarse-grained block structure became less pronounced. Other cell types showed inverted or reciprocal trends, but overall aging appears to involve a shift between the single gene and genome-wide ensemble scales.

## 3 Discussion

In this work, we proposed a physics-inspired framework to analyze high-dimensional single-cell transcriptomic data across multiple scales using coarse-graining methods inspired by the renormalization group. This approach views genome as interacting on a virtual grid and applies coarse-graining to construct an spectral language capturing subtle features not directly accessible at the single gene level. Tracking distribution evolution under coarse-graining integrates information across resolutions to uncover multiscale signatures. On the other hand, this method isn’t to explicitly define clusters like clustering methods or define a single-scale representation like dimensionality reduction. The coarse-graining flow is constructed in a successive way that information is gradually carrier over the joint probability distribution.

Then, we have demonstrated how the approaches can be used in analysis of gene expression data. Based on the established baseline Gaussian behavior, we proposed that two metrics can be of use to effectively quantify the coarse-graining flows. Our spectral analysis revealed distinct aging dynamics across cell types, likely reflecting their unique properties and functions. Further exploring the underlying mechanisms driving these differences can be a future direction. The proposed coarse-graining framework isn’t to define a single dimensionality or novelty but to incorporate the information from coarse-graining into a ‘spectrum’, which provides a more thorough analysis of aging dynamics in gene expression. For instance, in spleen B cells, the original scale lacked insight, but coarse-graining uncovered vanishing block structure indicative of increasing randomness. Conversely, limb muscle cells showed clear single-gene trends, but coarse-graining helped remove noisy modes to reveal true aging-related dynamics. Overall, jointly modeling the full coarse-graining spectrum gives a more thorough characterization of aging.

Next, the analysis expanded to more cell types suggest that it might not be optimal to seek for a single dimensionality while studying aging dynamics. Gene expressions of different ages can exhibit similar features at certain scale. However, the multiscale description revealed by coarse-graining isn’t to define a single dimensionality or novelty but to incorporate the information from coarse-graining into a ‘spectrum’, which provides a more thorough analysis of aging dynamics in gene expression. Finally, there was also one intriguing finding, the shift between normality and spectral gap. which may provide insight into future direction in aging studies.

Additionally, here we quantified flows using normality and spectral gaps, motivated by the establishe baseline bahavior. Intriguingly, these simple metrics revealed a shift between single-gene and genome-wide ‘randomness’ during aging. From the maximum entropy principle, decreased normality test statistics or Gaussian-like marginal distributions indicates increased single gene randomness. Conversely, decreased spectral gap or vanishing block structure indicates increased genome-wide randomness. Such a shift of ‘randomness’ was a notable universal signature from the multiscale coarse-graining analysis,. Aging may be better understood not as simply increasing randomness, but as a shift in randomness between scales. Extending this approach by characterizing additional flow features could further improve multiscale modeling of aging.

Moreover, the heart of conventional renormalization group is to see the fixed point or an independent operator. The apparent convergence to non-Gaussian distributions hints that simpler generative models may explain single-cell gene expression profiles at a given age. Learning such models from data could reveal underlying rules governing aging transcriptomics. Overall, this study demonstrates the value of coarse-graining for extracting biological insights from complex high-dimensional single-cell data in a physics-inspired manner.

## Supporting information

### S1 Appendix & Fig. Data Normalization

The normalization method used in this work is analytic Pearson residuals [15]. It is based on a common modeling assumption for count data without biological variability. Assume each gene *g* takes up a fraction *p*_*g*_ of the total amount *n*_*c*_ of counts in cell *c*. The read counts are then modeled as Poisson or negative binomial samples with expected value *μ*_*cg*_ = *p*_*g*_*n*_*c*_ without zero-inflation:

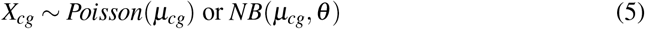

where *θ* is a dispersion parameter. The expected value of a gene count is then divided into two parts: gene-specific effect and cell-specific effect, *μ*_*cg*_ = *n*_*c*_ *p*_*g*_. In the Poisson model, the MLE solution is exact: 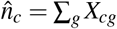 (sequencing depth) and 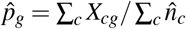. Or equivalently,

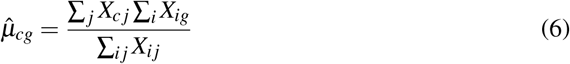

which proves to be a good approximation of MLE solution for negative binomial model. Then the normalized counts are given by:

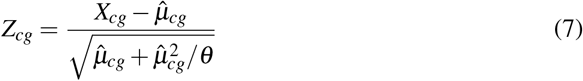

where the denominator 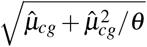 is the approximated NB variance. We use he most common practice of the choice with *θ* = 50.

This normalization method restores the gene counts into their analytical *z*-score, which is assumed to have zero mean and unit variance (Fig 6).

**Fig 6.**
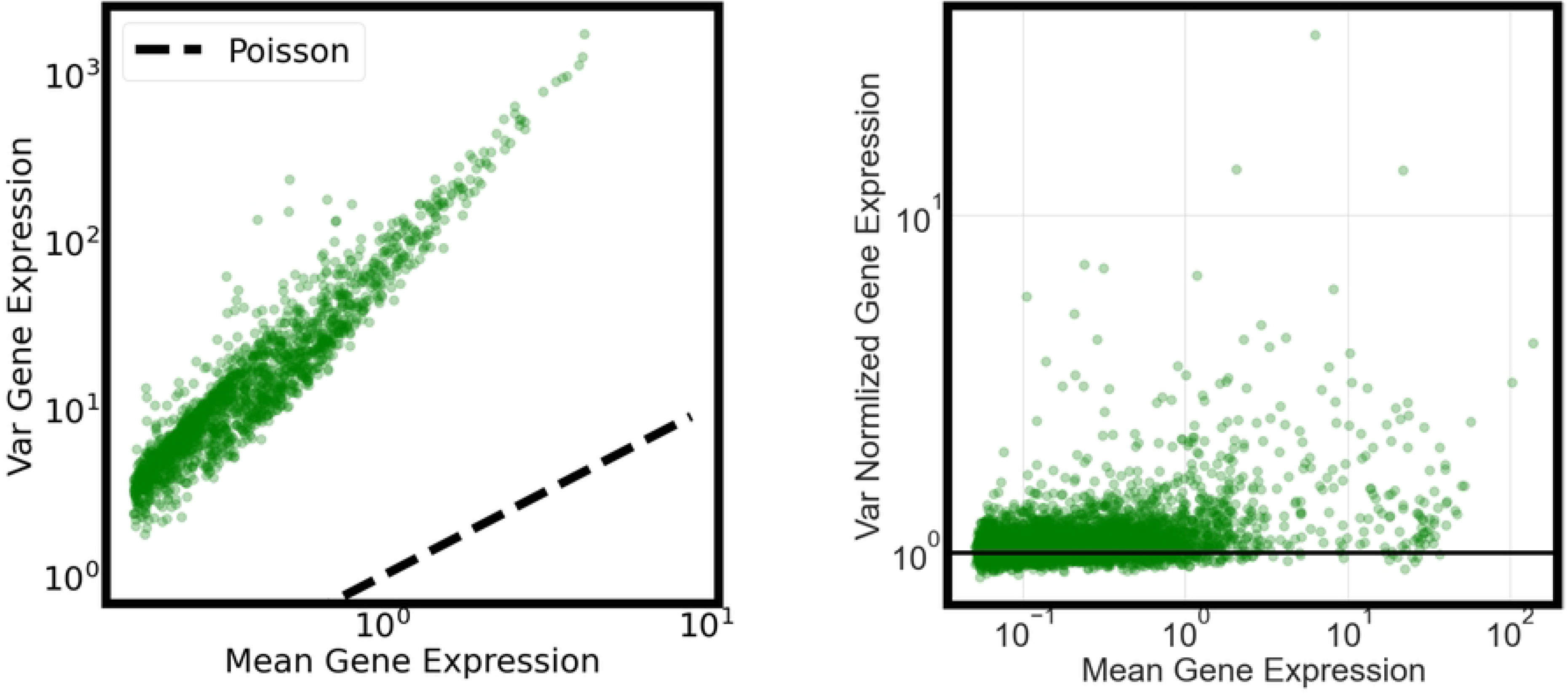
Pearson Analytical Residual. Left panel: the var-mean plot of the raw counts. Right panel: the var-mean plot of normalized gene expression.

### S2 Appendix. Quality Control

Before normalization, it is often necessary to perform a quality control on the raw data, in order for a better normalization performance. It is often done by screening out poorly sequenced genes. An empirical choice for Tabula Muris Senis it to screen out genes that are expressed in less than 5% of sequenced cells. This choice has two considerations, leftover genome size and normalization performance. First, it is not desirable to screen too many genes, which may hamstring the downstream analysis. At a threshold around 5%, we are about to keep 60% of the genome for all cell types. If the threshold goes up to 20%, the genome size will be overly reduced. On the other hand, in terms of variance stabilization, this threshold does yield normalized counts with the variance of gene counts being well stabilized around 1.

Also, when studying aging dynamics, it is necessary to study the same gene set that is aging. Only in this way can we ask to look into the progression of correlation structure. That said, for different age groups, the quality control should yield the same gene set. Only this way can we discuss the ‘aging’ of the entire genome. This is achieved by screening out the union of poorly read genes for three age groups simultaneously.

### S3 Appendix & Fig. Null Analysis

In the empirical setting, a null analysis seems appropriate. As prescribed in the main text (Sec 2.1), a null data is generated by marginal resampling. To be more specific, a gene expression data is an *N*-by*M* matrix, where *N* is the number of genes and *M* is the number of sequenced cell. By marginal resampling, we simply shuffle each row of the gene expression matrix. That is, the individual distribution of every single gene is preserved. However, by such a resampling, the correlation structures no longer exist, giving us an appropriate null model to investigate.

Fig 7 demonstrates both the coarse-graining methods applied to the null data. In panal (a.ii) and (b.ii), we can clearly see that both coarse-graining methods eventually converge to a standard normal distribution. (a.i) further confirms the fact. The block structure in correlation matrices would also never appear since any correlations have been destroyed by the marginal resampling. So the eigenvalue spectrum would be nearly continuous and flat as predicted by Marchenko-Pasture distribution and the corresponding spectral gap would be super small.

**Fig 7.**
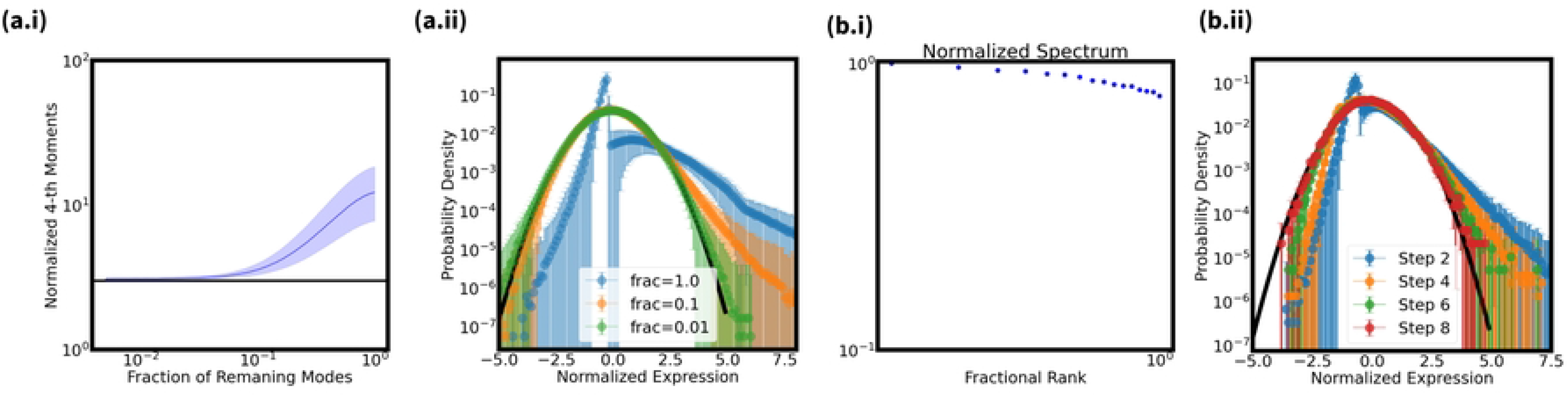
The coarse-graining analysis for marginal-resampled gene expression. **(a.i):** The normalized 4th order moments of momentum-space coarse-graining flow. **(a.ii):** The joint distribution of momentum-space coarse-graining flow. **(b.i):** The normalized eigenvalue spectrum of a marginal resampled matrix. **(b.ii):** The joint distribution of real-space coarse-graining flow.

### S4 Appendix. Usage of Quantitative Analysis

#### Spectral Gap

We used spectral gap to assess the block structure in correlation matrices. It can be easily related to the block structure by principal component analysis. In traditional PCA, a block in the correlation matrix will correspond to a large-variance principal component. Suppose two correlation matrices have similar block structures. The spectral gap among blocks or between the smallest block and other modes is maximized when all the blocks are uncorrelated. That being said, the block structures are outstanding. Otherwise, if there are many interwinding correlations among blocks, the spectral gap will shrink. Eventually, when the block structure vanishes, the eigenvalue spectrum is expected to be continuous so the spectral gap would be very small.

#### Anderson-Darling Test

The formal definition of the Anderson-Darling normality statistics [17] for an empirical set {*Y*_*i*_} is

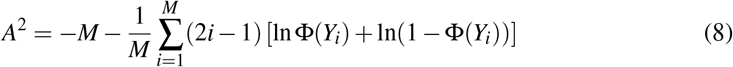

where Φ is the standard normal cumulative distribution function. As suggested by D’Agostino (1986), the statistics should be adjusted as

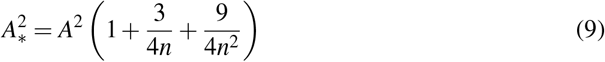

Apparently, the Anderson-Darling is associated with the sample size. In order to eliminate the effect of different sample size when we compare the normality across different groups, we performed bootstrap. More specifically, since the number of sequenced cells are 3,000 for each group, we use bootstrap to resample 6,000 cells for every single gene when calculating the Anderson-Darling normality statistics.

### S5 Appendix. Data Availibility

Tabula Muris Senis data is a high-through RNA-sequencing data of mice for different ages that was developed by Tabula Muris team. The data is currently available at figshare and more information can be found in the original paper: https://www.nature.com/articles/s41586-020-2496-1.

